# *Lactobacillus acidophilus* disrupts collaborative multispecies bile acid metabolism

**DOI:** 10.1101/296020

**Authors:** Sydney Dautel, Nymul Khan, Kristoffer R. Brandvold, Colin J. Brislawn, Janine Hutchison, Karl K. Weitz, Heino M. Heyman, Hyun-Seob Song, Zehra Esra Ilhan, Eric A. Hill, Joshua R. Hansen, Xueyun Zheng, Erin S. Baker, John R. Cort, Young-Mo Kim, Nancy G. Isern, John K. DiBaise, Rosa Krajmalnik-Brown, Janet K. Jansson, Aaron T. Wright, Thomas O. Metz, Hans C. Bernstein

## Abstract

Bile acids are metabolic links between hosts and their gut microbiomes, yet little is known about the roles they play in microbe-to-microbe interactions. Here we present a study designed to investigate the effect that a common probiotic, *Lactobacillus acidophilus*, has on microbial interactions that lead to formation of secondary bile acids. A model microbial consortium was built from three human gut isolates, *Clostridium scindens, Collinsella aerofaciens,* and *Blautia obeum*, and cultured under different bile acid and probiotic treatments. A multi-omics platform that included mass spectrometry-based metabolomics and activity-based proteomic probes was used to produce two major results. The first, was that an uncommon secondary bile acid – ursocholate – was produced by a multi-species chemical synthesis pathway. This result highlights a new microbe-to-microbe interaction mediated by bile acids. The second finding was that the probiotic strain, *L. acidophilus,* quenched the observed interactions and effectively halted consortial synthesis of ursocholate. Little is known about the role that ursocholate plays in human health and development. However, we did discover that a decrease in ursocholate abundance corresponded with successful weight loss in patients after gastric bypass surgery versus those who did not lose weight after surgery. Hence, this study uncovered basic knowledge that may aid future designs of custom probiotic therapies to combat obesity.

## INTRODUCTION

The human gastrointestinal (GI) tract is a complex ecosystem that functions in symbiosis with oral and intestinal microbiomes^1,2^. It has long been recognized that the composition of the gut microbiome has a significant effect on host digestion but more recent research has implicated the microbiome in human health and disease states that include cardiovascular disease risk^3^, neurological function^4^, and autoimmunity^5^. Rapid gains in knowledge of host-microbiome associations will undoubtedly lead to new practical applications^6^. Of these, the use of probiotics to modulate both the function and composition of gut microbiomes is especially promising^7,8^. Probiotic supplementation likely reduces the risk of developing antibiotic associated diarrhea^9^ and necrotizing enterocolitis in infants^10^. However, the therapeutic opportunities for probiotics are advancing to more precisely target specific processes carried out by the gut microbiome to impart health benefits beyond enhanced digestion^11-13^. Probiotic therapies are being explored relieve symptoms of autism^14^, depression^4^, autoimmune diseases^15,16^, and irritable bowel syndrome^17^ among many other conditions with positive – albeit conflicting – results. The efficacy of probiotic treatments is variable. Differing results can obviously arise from inconsistent study design – e.g., probiotic strain, dose – trial size, but they are also indicative of a large scientific knowledge gap and incomplete understanding of the mechanisms by which probiotics impact the GI-tract microbial ecosystem.

There are many hypotheses about the modes of action by which the gut microbiome and probiotic microbes impact human health. One proposed model is through modulation of the host immune system^18^, which has been concluded from studies that showed probiotic treatments affecting host immune function in humans facing pathogenic challenges^19-21^ or with autoimmune disorder^15,16,22^. Another hypothesis is that probiotics increase microbial competition within the intestinal ecosystem, thereby making it more difficult for pathogenic bacteria to survive^23,24^. Studies have also speculated that observed therapeutic action from probiotics results from their effect on the intestinal physiology by modulating endothelial junctions^25^ and the mucosal lining^12,25^ through a variety of proposed metabolic pathways. Another possible mode of action for probiotics is through bile acids. It has long been known that bile acids are important linkers between host and gut microbes. Intestinal bacteria produce secondary bile acids by deconjugation, reduction, oxidation, and epimerization of their host’s primary bile acids. Many probiotics can alter bile acid pools in humans^26,27^.

The primary bile acids (cholic acid and chenodeoxycholic acid) are synthesized in human hepatocytes and conjugated to the amino acids glycine and taurine to increase solubility^27,28^. They are then released into the duodenum and moved through the small intestine to assist emulsification of dietary lipophilic substances^29^. Approximately 95% of the bile acids are then passively reabsorbed in transit through the small intestine, resulting in approximately 5% (400-800 mg) passed on to the colon^27^. In the colon, these bile acids are rapidly deconjugated by the microbiome and reduced, oxidized and epimerized to a variety of secondary bile acids^27^. These secondary bile acids are known to have diverse effects on human health ranging from direct cytotoxicity^30^, to altered probability of cancer^31^, to hormonal function as cell messengers^29,32-34^. More recently, this list of known host-related effects has grown to include modulating the composition and function of the gut microbiome^35,36^ – e.g., by disassembling lipid membranes.

The intestinal microbiome represents a major modifier of the human bile acid pool. This is evinced by the fact that bile in the gall bladder is comprised of 70% primary bile acids but only 4% primary bile acids in the feces^27^. Not only does the microbiome determine bile acid composition, but bile acids also direct microbial communities^37^. For example, studies have shown that high fat diets and diets high in resistant starch^38^ have an effect on both the bile acid pool^39^ and the microbiome^40^. It has also been shown in rats that oral administration of certain bile acids can shift the microbiome composition^35,41^. Collectively, these studies suggest that there is a complex interplay between microbial species and bile acids that is not fully understood.

The relationships between the gut microbiome, probiotics and the bile acid pool are particularly relevant due to the current epidemic of obesity and the comorbidities associated with high adiposity (high cholesterol, high blood pressure, diabetes)^42^. Due to high rates of obesity, bariatric surgery has become a more common procedure with the number and types of surgeries increasing with time^43,44^. It is now well established that bariatric surgery results in reduced weight and a reduction in many comorbidities of obesity^45^. Several different advantages (weight loss, reduction of comorbidity) and disadvantages (surgical complications, malnutrition, weight regain, re-surgery, infection, etc.)^45,46^ have been have been identified for different bariatric procedures. Yet, there remains an incomplete understanding of the exact mechanisms of many of these outcomes, making it difficult to predict which patients will benefit most from these procedures. Elucidation of the root causes will require consideration of the impacts that the bile acid pool, microbiome composition and probiotic administration can have^47,48^. Deeper mechanistic insight could not only result in increased surgery success, potentially by pre-emptive modulation of the microbiome by bile acid pool^49,50^, but could also result in less surgeries necessitated if some of the positive results can indeed be realized via targeted use of probiotics.

Here we present a study that was designed to investigate the community dynamics of microbes commonly found in the gut and the impact that both the addition of bile acids and a probiotic has on interspecies interactions. An *in vitro* model was built as a three-species bacterial consortium, *Clostridium scindens, Collinsella aerofaciens,* and *Blautia obeum,* each of which occurs naturally in the human gut. This consortium was then treated with the probiotic strain *Lactobacillus acidophilus* to make an altered four-member consortium. Multi-omics assays were used to elucidate the interspecies microbial interactions with and without bile acid treatments (cholic acid and deoxycholic acid). We found that a secondary bile acid, ursocholic acid (7-epicholic acid), was produced from cholic acid through a multispecies chemical synthesis route mediated by an interaction between *B. obeum* and *C. aerofaciens*. This process was quenched by the addition of *L. acidophilus*, which disrupted the coordination between *B. obeum* and *C. aerofaciens* and shut down ursocholic acid production. These results were then contextualized by performing targeted metabolite measurements in fecal samples from a human clinical study^44^ that investigated secondary bile acid abundances as outcomes of gastric bypass surgery. The abundance of ursocholic acid corresponded with successful post-operative weight loss, highlighting that it may be important to investigate the implications of both patient- and microbe-derived metabolites to help gain a predictive understanding of a patient’s response to bariatric surgery. New knowledge in this area will yield opportunities to design custom probiotic therapies. More broadly, this study supports an emerging theme in microbiome sciences that microbial interactions are context dependent and the presence or absence of select species and/or metabolites can have a strong effect on the overall function.

## RESULTS

### A model microbial consortium responds to probiotic and bile acid treatments

We constructed an *in vitro* microbial consortium from human gut bacterial isolates. A 3-member consortium (3MC) composed of *Clostridium scindens* ATCC 35704*, Collinsella aerofaciens* ATCC 25986, and *Blautia obeum* ATCC 29174 was compared to a 4-member consortium (3MC + L) that included an added a probiotic strain, *Lactobacillus acidophilus* ATCC 4356. The species were chosen for this model based on their capacity to perform one-of-three unique microbial transformations on human bile acids (Table S1): 7α-dehydroxylation, hydroxysteroid dehydrogenatation, or bile salt hydrolsis^27,37,51,52^.

Anaerobic culturing treatments included supplementation with cholic acid and/or deoxycholic acid and were compared to axenic controls. We found that each member of the consortia grew together under anaerobic culturing conditions but that each consortium (3MC versus 3MC + L) had very different growth and extracellular metabolite profiles (Fig. 1). Despite the anticipated anti-microbial effect of bile acids^26,53,54^, all cultures in media containing added cholic acid and deoxycholic acid (0.1 mM) showed faster specific growth rates compared to their corresponding treatments without bile acids (Fig. 1a and b). The axenic controls confirmed that *L. acidophilus* had the fastest specific growth rate, followed by *B. obeum*, *C. aerofaciens* and *C. scindens*. The probiotic, *L. acidophilus*, was the least affected by the addition of the bile acids to the growth media, with only a 17% increase in the specific growth rate as compared to 170%, 76% and 167% increases for *C. scindens*, *C. aerofaciens* and *B. obeum,* respectively. An adonis test was used to show that the species composition of each consortium – i.e., presence/absence of *L. acidophilus* – was the strongest determiner of variance (R2 = 0.49, p < 0.001) in the extracellular metabolome. This was in contrast to the effect of treatments that tested for changes in the global metabolome based on bile acid or no bile acid inputs, which were not a statistically significant source of variance (Fig. 1C). Hence, the probiotic *L. acidophilus* was a major modifier of the extracellular chemical environment.

**Figure 1.**
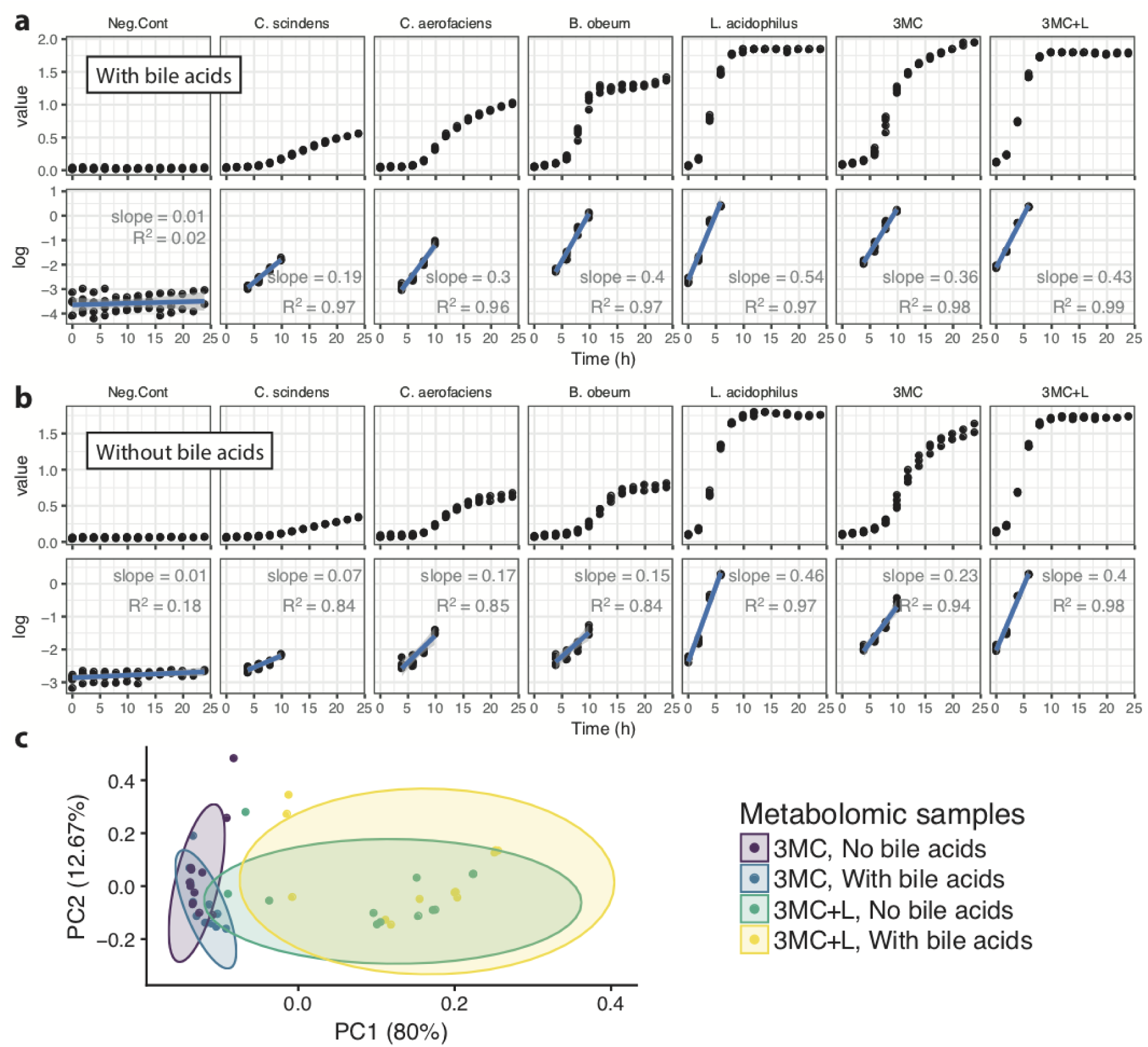
Probiotic influence on consortial growth and the extracellular chemical environment. (A) Growth of the 3-member consortium (3MC) in bile acid supplemented media as compared to (B) the 4-member consortium (3MC + L) that differed by the addition of probiotic, *L. acidophilus.* The y-axis of A and B represent means from triplicate measurements of absolute species abundance defined as the total optical density (OD_630nm_) at each time point multiplied by each respective measurement of relative abundance obtained from qPCR; error bars represent ± 1 standard deviation. (C) A principle component analysis on the extracellular metabolome ordinated by the Euclidean distance between the metabolic profiles from each treatment. The colored ellipses represent 95% confidence limits assuming a multi-variate t-distribution.

### *L. acidophilus* quenches secondary bile acid production

Ursocholic acid was produced in the 3MC (Fig. 2A), but not by any one species grown under axenic conditions. Hence, ursocholic acid was produced by a multispecies synthesis route that required at least two species from our model consortium. Addition of the probiotic, *L. acidophilus* quenched the production of ursocholic acid to negligible levels as compared to those measured in the 3MC (Fig. 2b). In addition to ursocholic acid production, *L. acidophilus* also attenuated growth of *B. obeum* and *C. aerofaciens* as observed in the 3MC + L species-specific growth dynamics, which were in stark contrast to the 3MC (Fig.2c). Within the 3MC + L, *L. acidophilus* became a dominant member of the community, but in the absence of the probiotic, the 3MC was dominated by *B. obeum* with *C. scindens* showing the lowest relative abundance in both consortia.

**Figure 2.**
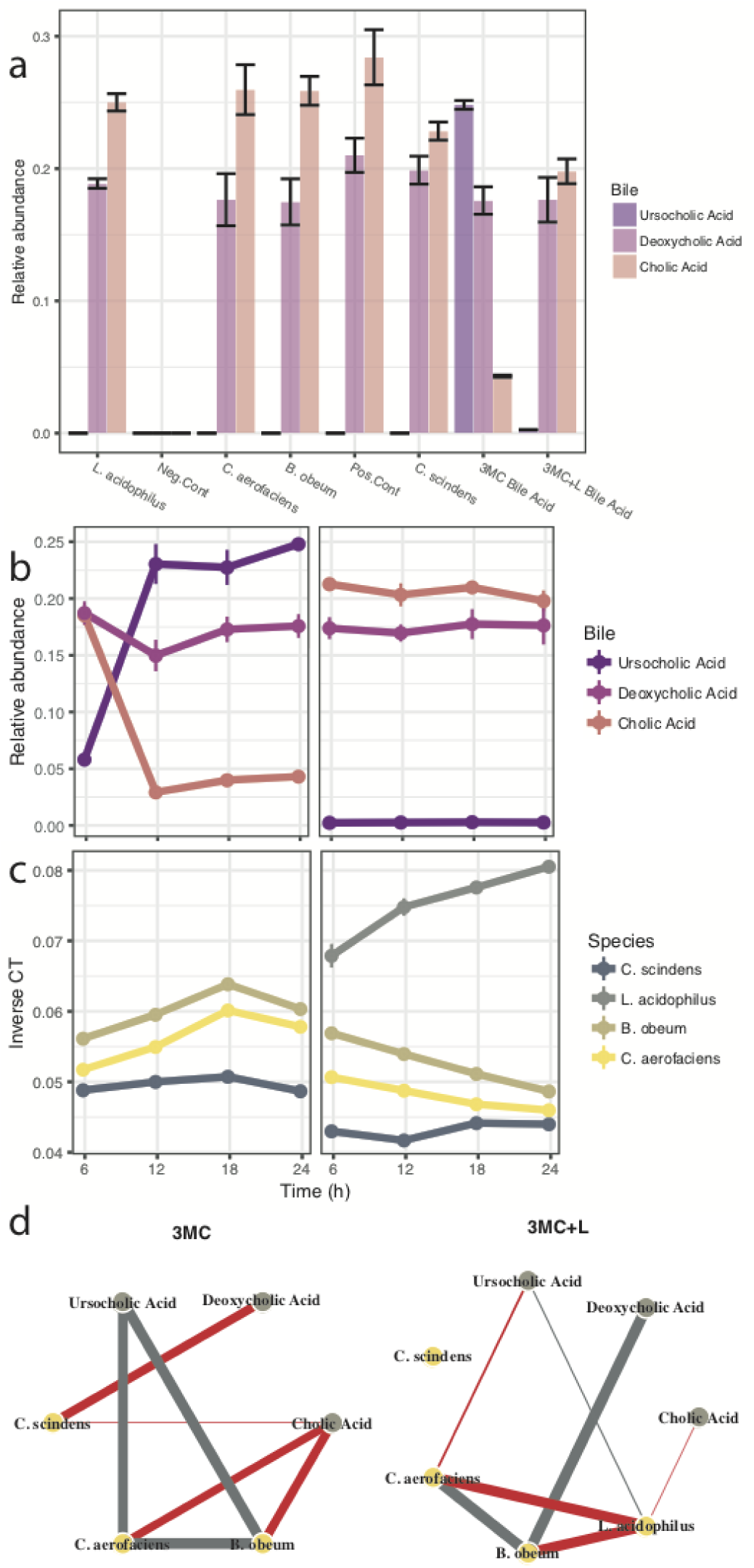
Secondary bile acid – ursocholic acid – was produced by consortia but not axenic cultures; *L. acidophilus* disrupted the consortial synthesis. (A) Comparative abundances of bile acids measured after 24 h incubations (3MC and 3MC + L) and axenic controls. Time course measurements of (B) bile acid abundances and (C) species abundance as shown by the inverse qPCR cycle thresholds (CT). Comparisons are shown between the 3MC and 3MC + L (with *L. acidophilus*) treated with the 0.1 mM bile acid mixture. Each data point shown in panels A-C represent the mean from a minimum of three biological replicates ± 1 standard deviation. (D) Pearson’s correlations between bacterial species and bile acid abundances; thicker lines correspond to greater correlation coefficients (cut-off below 0.85); red and grey colors correspond to positive and negative correlations, respectively.

The dynamic profiles of bile acids and bacterial species were correlated for each treatment group, 3MC and 3MC + L (Fig. 2d), respectively. Pearson’s correlations were determined between species and for species to bile acids but not between bile acids. The results show that ursocholic acid shared strong positive correlations (Pearson’s coefficient; r > 0.85) with the abundances of *C. aerofaciens* and *B. obeum* in the 3MC. As expected, cholic acid showed strong negative correlations with *C. aerofaciens* and *B. obeum* within the 3MC, establishing that it was the most likely substrate for ursocholic acid synthesis. Cholic acid, was not correlated with *C. scindens*, implying that this species may not have been involved in production of secondary bile acids directly from cholic acid. The lack of *C. scindens’* participation in secondary bile acid synthesis in the 3MC was also evinced by its strong negative correlation with deoxycholic acid, a known 7α-dehydroxylation product of *C. scindens* that utilizes cholic acid as the substrate^27^.

*L. acidophilus* membership changed the correlations between species and bile acids. Most notably, ursocholic acid was not correlated with any species in the 3MC + L under the probiotic treatment. *L. acidophilus* did not correlate with any of the bile acids and shared strong negative correlations with *C. aerofaciens* and *B. obeum*, indicating that competition and/or antagonism are the likely mechanisms by which the multi-species ursocholic acid synthesis was quenched.

### Multispecies synthesis of ursocholic acid

Based on the metabolomics results, we hypothesized a multi-species cooperative synthesis of ursocholic acid that involved both *B. obeum* and *C. aerofaciens*. The initial inference was derived by comparing the metabolite profiles between the axenic and consortial treatments (Fig. 2a). The next piece of evidence was obtained from correlations between species and bile acids in the context of *a priori* knowledge of the metabolic reactions that were the basis for choosing each species in the model consortium (Fig. 2d). A possible mechanism for this could start with *B. obeum* conversion of cholic acid into a transient 7-keto intermediate, such as 7-oxodeoxycholic acid. A ketone intermediate was not identified by our bile-acid-targeting LC-MS metabolomics approach but that does not exclude the possibility of its existence. The next step could be achieved by the known genome encoded functions of *C. aerofaciens*, which contains *hdhB* (GenBank accession ZP_01773061)^55^. This gene encodes for a 7β-hydroxysteroid dehydrogenase (7β-HSDH), known to catalyze a reaction that takes a 7-oxodeoxycholic acid to ursocholic acid^56^. However, the *B. obeum* ATCC 29174 genome does not contain an oxidoreductase that is clearly annotated to catalyze our hypothesized reaction in this first step. *B. obeum* does contain a *baiA* gene (RUMOBE_03494) that encodes a putative 7α-dehydroxylase, but genes encoding for a 7α-HSDH have yet to be identified. Yet our experiments clearly showed synthesis of ursocholic acid and we confirmed a required participation of *B. obeum*; hence, we concluded that a non-specific or previously unidentified oxidoreductase catalyzed this first step.

To test our hypothesis that *B. obeum* expresses enzymes that react with cholic acid (other than the known BaiA protien), we synthesized and employed a custom cholic acid photo affinity probe (CAP; cholic acid probe). This probe is a cholic acid derivative designed to bind and enrich proteins that can then be identified by MS-based proteomics (Fig. 3a). The results were a suite of proteins from each species in the consortium that were significantly enriched by the CAP (>10^2^ fold-change; p-value < 0.001) (Table S2). Surprisingly, the *B. obeum* BaiA protein (*RUMOBE_03694*) was not enriched by the CAP, indicating the possibility of an incorrect annotation from homology of UniProt-KB A5ZWVO or lack of expression in these consortial conditions. We identified a suite of *B. obeum* proteins that were significantly enriched by the CAP and specific to the treatments corresponding to ursocholic acid synthesis and the *B. obeum* axenic treatment. Of these, we focused on a pair of oxidoreductases as candidates for the hypothesized two-step, two-species reaction that lead to ursocholic acid in our consortium. While they did not share strong homology with known 7α-HSDH proteins, they were annotated as enzymes that may catalyze the hypothesized alcohol-ketone inter-conversions (Fig. 3b). These CAP-binding proteins were annotated based on homology as an alcohol dehydrogenase (A5ZM66; *RUMOBE_00083*) and an alcohol-aldehyde dehydrogenase (A5ZNA3; *RUMOBE_00470*).

**Figure 3.**
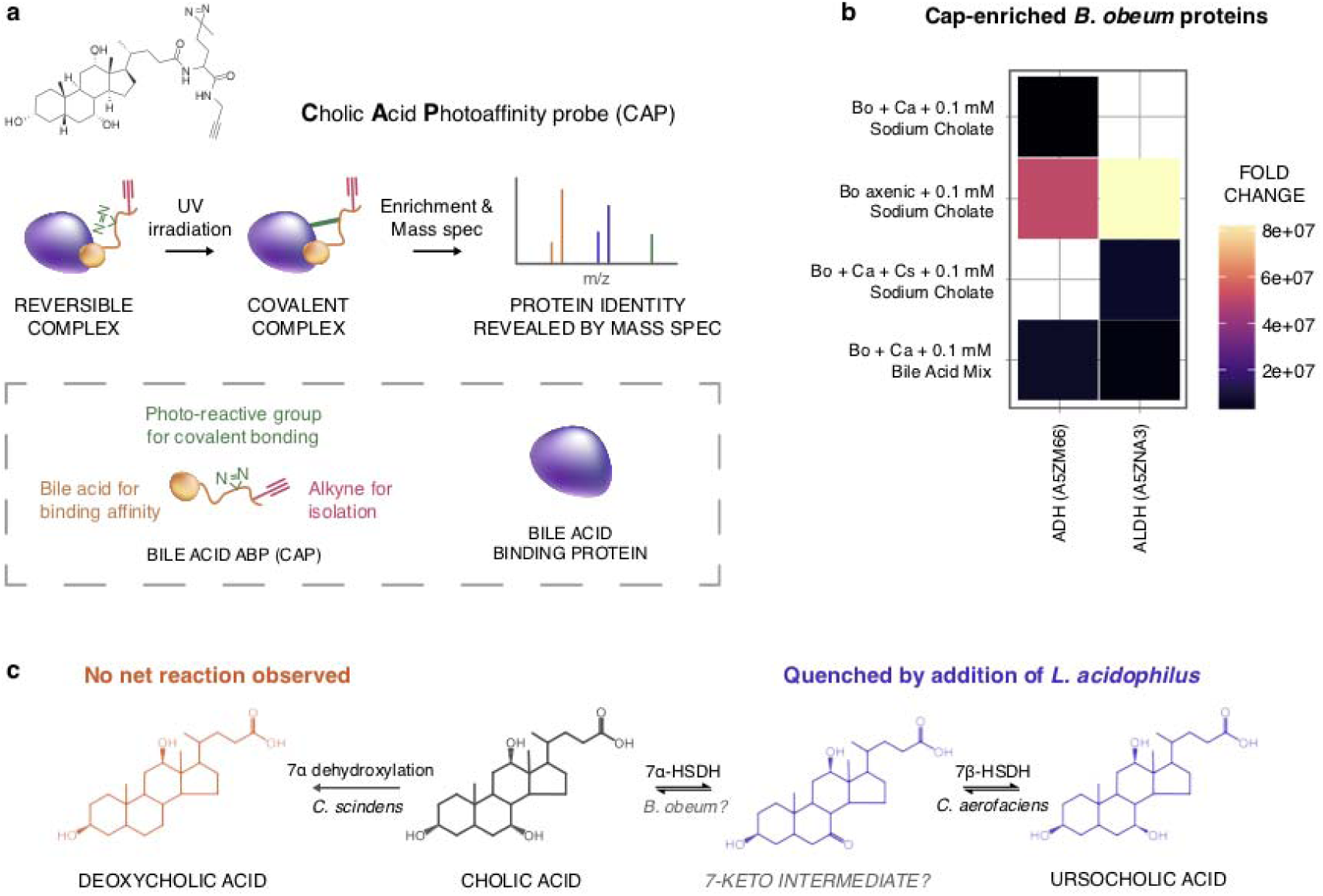
The multispecies synthesis hypothesis and cholic acid binding oxidoreductases. (A) The cholic acid photoaffinity probe (CAP) was synthesized and used to enrich proteins for mass spectrometry-based proteomics. (B) Of the proteins enriched by CAP (> 100 fold-change and p < 0.001), we identified two oxidoreductases annotated alcohol or alcohol-ketone dehydrogenases (ADH and ALDH, respectively). These proteins are B. obeum candidates for the hypothesized two-step, two-species mechanism (C) leading to the synthesis of ursocholic acid. The abbreviations used to describe experimental treatments are as follows: Bo (*B. obeum*); Ca (*C. aerofaciens*); and Cs (*C. scindens*).

### Implications toward obesity and bariatric surgery

The proposed mechanism for physiological effect for Roux-en-Y gastric bypass surgery (RYGB) are BRAVE “Bile flow alteration, Reduction of gastric size Anatomical gut re-arrangement, Vagal manipulation, Enteric gut hormone modulation^57^. Hence, the GI tract of patients that have undergone bariatric surgery is a potential model system to study bile acid metabolism/alterations given that bile acid profiles change by increasing abundance of secondary bile acids^49^. To investigate the clinical relevance of ursocholic acid, we leveraged access to a cohort of patients that had undergone gastric bypass surgery. Targeted measurements of ursocholic acid were performed and compared in fecal samples collected from 24 patients that underwent RYGB surgery: 10 patients with normal weight and 14 morbidly obese controls, which included those scheduled for surgery^44^. We found that the abundance of ursocholic acid corresponded with obesity and the success of gastric bypass surgical procedures (Fig. 4). Success was defined when patients exhibited at least 50% excess weight loss and less than 20% regain. Ursocholic acid levels were significantly higher in the morbidly obese controls (pre-surgery) compared to patients that had experienced successful gastric bypass surgery. There was no statistically significant difference between the fecal abundance of ursocholic acid in the obese controls and the unsuccessful surgical patients.

**Figure 4.**
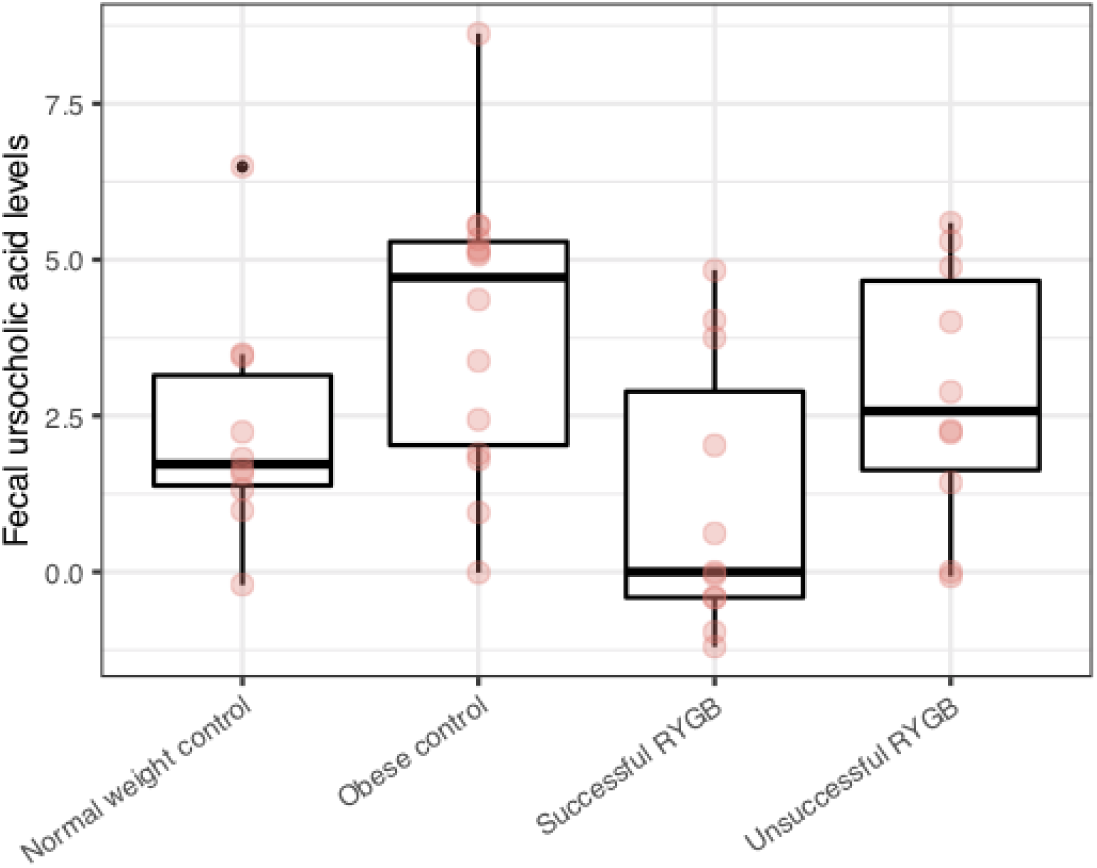
Normalized abundance of ursocholic acid in feces of patients who had undergone gastric bypass surgery – Roux-en-Y gastric bypass (RYGB) – as compared to normal weight and obese controls. Unpaired two-tailed t-tests were used to infer a statistical difference between the means of the obese controls – a group that contained pre-operative patients – and those that experience successful RYGB (p < 0.005). Different ursocholic acid abundances were also observed – albeit statistically less conclusive – between successful and unsuccessful RYGB patients (p < 0.076).

## DISCUSSION

The human gut microbiome is integral to human health and development^58,59^. However, the function and composition of the human GI-tract ecosystem is complex^60^, which often makes detailed studies of individual processes difficult. The use of model systems is a valuable approach to dissect complex biological functions. In particular, simplified model consortia gaining interest in microbiome research because they enable experimentalists to manage complexity by controlling multiple system components under defined treatments. The utility of simplified consortia, or bench-top microbiomes, has been demonstrated for a variety of human-associated communities^61,62^ and other complex microbial ecosystems related to plants^63^, sediment/biofilms^64-66^ and marine habitats^67,68^. Here, we developed a simple model microbial consortium that was specifically designed to investigate how the addition of a common probiotic (*L. acidophilus*) effects microbial interactions mediated through bile acid transformations. This study was not designed to directly inform microbial physiologies that should occur in the human GI-tract *in situ*. Rather, it was designed and successfully implemented for controlling the membership of microbial species and presence/absence of bile acids that are associated with human digestive systems. Our model bile acid consortium helped produce two major results. The first was that ursocholic acid was synthesized by the coordinated metabolism of a consortium and not by any single species included within this study. The second important finding was that probiotic, *L. acidophilus* quenched the observed multi-species interactions that resulted in secondary bile acid synthesis.

Ursocholic acid is the 7 beta-hydroxyepimer of cholic acid. It is rarely mentioned in the scientific literature and has been termed an “unusual secondary bile acid” as compared to more commonly studied metabolites such as deoxycholic and ursodeoxycholic acid^69^. Previous studies have investigated ursocholic acid as a potential therapeutic to modulate the host’s synthesis of primary bile acids^70^, or to improve the liver function of patients with primary biliary cirrhosis^71^ and reduction of bile cholesterol saturation^72^. In these previous studies, ursocholic acid was typically tested along with ursodeoxycholic acid and found to be notably less effective as a treatment for biliary cirrhosis^71^. Despite these therapeutic studies, little is known about the role that ursocholic acid plays in modulating human-microbe and/or microbe-microbe interactions.

The data derived from our clinical study showed that ursocholic acid is in fact present and abundant within the human GI-tract and its relative abundances change when drastic changes to microbiome occur (pre- and post-bariatric surgery)^44^. Our limited understanding of the role that ursocholic acid plays in human health and specifically the host-microbiome interactions that lead to its production represent a major knowledge gap. This is punctuated by the fact that our current study – and a previous study^73^ – have observed microbial synthesis of ursocholic acid and/or 7-oxodeoxycholic acid but not conclusively identified the genes and 7α-HSDH proteins responsible.

Cataloguing the bacterial genes from the “sterolbiome” is an active area of research^74^ that can yield new biological insight and help to improve human health by understanding how microbes modify chemical environments within the human GI tract. In this study, we hypothesized a multispecies chemical synthesis route in which *B. obeum* converts cholic acid into an intermediate ketone via a 7α-HSDH-like reaction which is then proceeded by the known 7β-HSDH reaction catalyzed by *C. aerofaciens*. Intraspecies 7-epimerization has been demonstrated in *Clostridium limosum*^75^ and *Clostridium absonum*^76^, which express both the required enzymes, 7α-HSDH and 7β-HSDH. However, genes encoding for 7α-HSDH, have yet to be identified in *B. obeum* and other bacteria such as an *Eggerthella* sp. known to express this protien^73^. Yet, we conclusively found that a collaborative reaction between *B. obeum* and *C. aerofaciens* does occur, which highlights an increase in our understanding of bile acid metabolism of bacteria.

We hypothesized a 7-keto intermediate that was transferred between *B. obeum* and *C. aerofaciens* in the observed multispecies chemical synthesis route. We did not identify an intermediate such as 7-oxodeoxycholic acid in the supernatant of the samples and therefore cannot categorically confirm its existence. However, 7α-HSDH mediated synthesis of 7-oxodeoxycholic acid has been previously observed in similar multi-step bile acid transformation processes^56^. It is possible that *C. aerofaciens* has a high affinity uptake mechanism for the hypothesized 7-keto intermediate such that extracellular concentrations were below the detection limits of our LC- and GC-MS metabolite identification methods. Another point of uncertainty is whether the CAP-enriched alcohol dehydrogenase (A5ZM66; *RUMOBE_00083*) and alcohol-aldehyde dehydrogenase (A5ZNA3; *RUMOBE_00470*) were responsible for the hypothesized reactions. Certainly, other proteins were enriched by CAP (Table S2), yet these were the only *B. obeum* proteins annotated with an enzymatic function capable of alcohol-aldehyde interconversion. We cannot rule out the possibility that enriched proteins of unknown function participated in the observed reaction. However, there is some precedent for associating secondary bile acid synthesis genes with alcohol dehydrogenases. BaiA proteins from *C. scindens*, encoding for 3α-HSDH proteins, have previously been shown to align well to short chain alcohol dehydrogenases in *Eubacterium* sp. Strain VPI 12708 and to alcohol/polyol dehydrogenase genes^77,78^. We chose to use *C. scindens* in the 3MC because it has genes that encode for Bai proteins and express HSDH proteins. In fact, the *C. scindens* reference proteome (VPI 12708) does contain a putative 7α-HSDH (UniProtKB – Q03906); however, *C. scindens* abundances and correlation-based inferences from this study did not provide evidence of *C. scindens’* participation in the transformation of cholic acid to secondary acids. Our conclusion was that *C. scindens* played a minor role in the system and was largely outcompeted by other members.

### Towards precision probiotics to complement bariatric surgery outcomes

The effect of probiotics on the human gut microbiome and the bile acid pools are particularly relevant due to the current epidemic of obesity and the comorbidities associated with high adiposity (high cholesterol, high blood pressure, diabetes)^42^. Hence, bariatric surgery is becoming more common as a treatment strategy. Yet knowledge gaps still exist with respect to how metabolites and microbes could play a role in successful outcomes. Here, we show that the abundance of ursocholic acid corresponds with the efficacy of gastric bypass surgery. This new knowledge about the role of specific probiotic strains and/or their metabolic products, are therefore leading towards promising novel treatments for patients undergoing bariatric surgery. It has already been shown that post-operative administration of *Lactobacillus* sp. improves weight loss and vitamin B absorption in RYGB patients^48^. It is possible that the cessation of ursocholic acid production or increased abundances of *Lactobacillus* sp. could result in better control over weight loss. We also note that there is some precedent derived from mouse models for the idea that probiotics or introduction of non-adapted microbial taxa can modulate a hosts’ microbiome^7^ and microbe-associated bile acid pool^79^.

### Conclusion

This current study was a fundamental investigation of microbial interactions and the role that a probiotic bacterium plays in modulating the synthesis of secondary bile acids. It was not intended to inform clinical practice. However, the results and conclusions presented establish an important idea related to broader microbiome sciences; microbial interactions are context dependent^64,65^ and the presence or absence of select species and/or metabolites can have a strong effect on the overall community-level function.

## METHODS

### Bacterial strains and cultivation

*Clostridium scindens* ATCC 35704, *Colllineslla aerofaciens* ATCC 25986, *Blautia obeum* ATCC 29174 (formerly *Ruminococcus obeum*) and *Lactobacillus acidophilus* ATCC 4356 were grown under axenic conditions and in consortia on Lactobacilli MRS Broth (BD Difco, Houston, TX, USA). The primary treatment the addition of bile salts, equivalent mixtures of cholic and deoxycholic acid (Sigma-Aldrich 48305, St. Louis, MO, USA), supplemented to 0.1 mM. The secondary treatment was the presence and absence of probiotic, *Lactobacillus acidophilus* ATCC 4356 rendering either a 3-member (*L. acidophilus* negative) or 4-member (*L*. *acidophilus* positive) consortia. Anaerobic growth conditions were prepared by boiling the media and subsequently sparging with an 80% N_2_, 10% H_2_, 10% CO_2_ gas mixture and transferring to 30 ml sealed Balch tubes under the same gas headspace prior to autoclaving. Each culture was inoculated to a starting OD_630nm_ = 0.077 ± 0.033 by each axenic cell suspension resulting in 1 serial passage of log phase cells; consortia were inoculated with an equivalent volume ratio mL. The optical density (OD_630nm_) was measured over a 24 h period using a Spectronic 20D+ spectrophotometer (ThermoSpectronic, Madison, WI, USA); each time point was sampled in triplicate via destructive sampling.

### PCR quantification of gene target

Bacterial DNA was extracted from consortia and axenic culture using the MoBio (Carlsbad, CA, USA) PowerSoil DNA Isolation Kit following the manufacturer’s protocol. A total volume of 5 µL of undiluted consortia DNA or standard curve DNA was analyzed in triplicate with an Applied Biosystems 7500 fast instrument (Foster City, CA). Samples were analyzed in triplicate with the primers shown in Table S3 targeting *rpoB*. PCR reactions were run using the FAST cycling conditions: initial denaturation was done for 20 seconds at 95 °C followed by 40 cycles of denaturation (95 °C for 3 seconds), annealing (60 °C for 30 seconds). The output from the real-time PCR assays were C_T_ values that represent the PCR cycle at which the amplification crosses a given threshold (0.1). All C_T_ values in Table S4; data are plotted and analyzed as inverse C_T_ representing the relevant abundance^66,80^ assuming equal between species in the consortia.

### Bile acid identification and quantification

Stock solutions of the cholic and deoxycholic acids (Steraloids Inc., Newport, RI, USA) were made in methanol (1 mg mL^-1^) and were then pooled together and diluted in a series in 0.1% formic acid to generate a 7 pt of the calibration curve (0.0062, 0.025, 0.050, 0.1, 0.5 and 1 μg mL^-1^). An internal standard (23-nor-5β-cholanic acid-3α, 12α-diol) was added to the filtered media (0.2 µm) collected from each microbial growth sample. Cold methanol (−20°C) was added at a ratio of 1:4 (filtrate:MeOH). The samples were mixed via vortexing, chilled at -20°C for 30 minutes and separated via centrifugation (1725 rpm, 10 minutes). The supernatant was removed and dried and then re-suspended in 0.1% formic acid in deionized water solution. These samples were analyzed on a Waters nano-Acquity UPLC system (Milford, MA, USA) configured for direct 5 µL sample injections onto an in-house packed fused silica column (360 µm o.d. × 150µm i.d. × 30 cm long; Polymicro Technologies Inc., Phoenix, AZ, USA) containing Waters HSS T3 media (1.8 μm particle size). A flow of 600 nL min^-1^ was maintained using mobile phases consisting of (A) 0.1% formic acid in water and (B) 0.12 % formic acid and 5 mM ammonium acetate in methanol with the following gradient profile (min, %B): 0, 1; 5, 1; 10, 65; 59, 99; 60, 1. Total run time including column re-equilibration was 75 min. Mass spectrometry (MS) analysis was performed using an Agilent model 6490 triple quadrupole mass spectrometer (Agilent Technologies, Santa Clara, CA, USA) outfitted with a custom nano-electrospray ionization interface built using 150 um o.d. × 20 um i.d. chemically etched fused silica^81^. The hexabore ion transfer tube temperature and spray voltage were held at 200°C and -4.0 kV, respectively. Data were acquired in negative ion mode for 75 min from sample injection using a dwell time of 200 µs, fragmentation of 380 volts, and collision energy of 10 volts. Selected reaction monitoring (SRM) transitions were acquired as shown in supplementary Table S5. Ursocholic acid was identified as an unknown in the initial LC-MS trials. After fractionation and purification, we isolated the unknown and verified that it was ursocholic acid via NMR and ion mobility mass spectrometry analyses. Authentic ursocholic acid was purchased from Toronto Research Chemicals (N. York, Ontario, Canada). Comprehensive details of these procedures are provided in the supplementary materials.

### Untargeted Metabolomics

The spent media was dried, chemically derivated and analyzed by GC-MS as previously reported^82^. GC-MS raw data files collected by GC-MS were processed using the Metabolite Detector software, version 2.5 beta^83^. Agilent .D files were converted to netCDF format using Agilent Chemstation (Agilent, Santa Clara, CA, USA) and then converted to binary files using Metabolite Detector. Samples were aligned chromatographically across all analyses after deconvolution. Metabolites were identified by matching experimental spectra to a Pacific Northwest National Laboratory (PNNL) augmented version of FiehnLib^84^. This library has spectra and validated retention indices for over 850 metabolites. In order to minimize errors in deconvolution and identification, all metabolite identifications were manually validated after automated data-processing.

### Bile acid photoaffinity probes and proteomics

Custom photoaffinity probes were synthesized as derivatives of cholic and deoxycholic acid for this study as described in detail within the supplementary materials. Bacterial lysate samples were normalized to 500 µL 1.8 mg/mL proteome in PBS buffer. Cholic acid photoaffinity probe (CAP) or an equal volume of DMSO control was incubated with proteome for 60 min at 37 °C. Final DMSO concentration was 1%. Samples were exposed to UV light (wavelength: 365nm; 115V, 15W) using a Fisher UVP95 lamp (Fisher Scientific, Hampton, NH, USA) for 7 minutes on ice. Subsequent to UV irradiation, the samples were subjected to click chemistry, with final concentrations of reagents being: biotin-azide (60 µM) in DMSO, sodium ascorbate (10 mM), THPTA (4 mM), and CuSO_4_ (8 mM). Each reagent was added individually in that sequence, vortexed, centrifuged, and incubated at room temperature in the dark for 90 min. 800 µL of pre-chilled MeOH was then added to each sample and incubated at -80 °C freezer for 30 min to induce protein precipitation. Samples were centrifuged at 14,000 × g at 4°C for 5 min. The supernatant was discarded, and the pellet was allowed to air-dry for 5 min. Samples were reconstituted and sonicated in 520 µl SDS (1.2%) in PBS; followed by incubation at 95 °C for 2 min. Samples were centrifuged at 14,000 × g for 4 min at room temperature. Protein concentrations were determined via BCA assay and samples were normalized to a volume of 500 µL at 1.2 mg/mL. Trypsin digestion was performed on protein bound to 100 µL Streptavidin-agarose beads. Peptides were reconstituted by adding 40 µl of 25 mM NH_4_HCO_3_ and heating the samples at 37°C for 5 min. Samples were transferred to ultracentrifuge tubes and were centrifuged at 100,000 × g to remove debris. 25 µL was added to glass vials for storage at -20 °C until analysis. All proteomics samples prepared for LC-MS were analyzed using a Velos Orbitrap MS as previously described^85,86^.

### Clinical data and experimentation

Fecal samples were collected from patients at the Mayo Clinic, Scottsdale, AZ, USA. The metabolomics assays were performed at the Pacific Northwest National Laboratory by the methods described above. The experimental design has been previously described^44^ and was approved by The Institutional Review Boards of Mayo Clinic and Arizona State University (IRB# 10-008725).

### Statistics

Analysis and graphing was performed in R^87^ making use of the ‘vegan’^88^ and ‘igraph’^89^ packages, along with many packages in the Tidyverse^90^. The adonis test (permutation MANOVA) was used to partition a matrix of Euclidean distances between global metabolite samples based on bile acid and probiotic treatments. An unpaired two-tailed Student’s t-test was used to compare fecal ursocholic acid measurements between treatment groups. Linear models were fit to the log of the exponential growth phase of microbial consortia, and for each regression, the goodness of fit and the probability of observing a similar slope if the true slope coefficient was zero was reported. Global proteomics was used to identify proteins enriched by the CAP probes, by selecting proteins with a fold-change increase of >100 and a t-test p-value of < 0.001. The methods for all hypothesis testing and descriptive statistical procedures are included in the R markdown supplied for this study.

### Data Repositories and Reproducibility

The raw data sets for this study along with the R scripts used for analysis and graphing are available from the Open Science Framework (OSF) under the name “Bile Acids Consortia” at https://osf.io/5meyd/. The mass spectrometry proteomics data have been deposited to ProteomeXchange with the dataset identifier PXD008617.

## ACKNOWLEDGMENT

This research was supported by the Microbiomes in Transition (M*in*T) Initiative, a Laboratory Directed Research and Development (LDRD) Program of Pacific Northwest National Laboratory (PNNL). Measurements of bile acids in samples from the human clinical trial were supported by the PNNL LDRD Program via the Signature Discovery Initiative. Research reported in this publication was supported by the National Institute of Diabetes and Digestive and Kidney Diseases of the National Institutes of Health under Award Number R01DK090379. The content is solely the responsibility of the authors and does not necessarily represent the official views of the National Institutes of Health. Metabolomics measurements and chemical characterization of ursocholic acid were performed in the Environmental Molecular Sciences Laboratory (user project 49356), a national scientific user facility sponsored by the Department of Energy Office of Biological and Environmental Research, located at PNNL. PNNL is operated for the DOE by Battelle under contract no. DE-AC05-76RLO-1830. Specific acknowledgments are given to Heather Brewer, Athena Schepmoes, Rose Perry and Erika Zink for their technical contributions.

